# Ionpair-π interactions favor cell penetration of arginine/tryptophan-rich cell-penetrating peptides

**DOI:** 10.1101/717207

**Authors:** Astrid Walrant, Antonio Bauzá, Claudia Girardet, Isabel D. Alves, Sophie Lecomte, Françoise Illien, Sébastien Cardon, Natpasit Chaianantakul, Manjula Pallerla, Fabienne Burlina, Antonio Frontera, Sandrine Sagan

## Abstract

Cell-penetrating peptides (CPPs) internalization can occur both by endocytosis and direct translocation through the cell membrane. These different entry routes suggest that molecular partners at the plasma membrane, phospholipids or glycosaminoglycans (GAGs), bind CPPs with different affinity or selectivity. The analysis of sequence-dependent interactions of CPPs with lipids and GAGs should lead to a better understanding of the molecular mechanisms underlying their internalization. CPPs are short sequences generally containing a high number of basic arginines and lysines and sometimes aromatic residues, in particular tryptophans. Tryptophans are crucial residues in membrane-active peptides, because they are important for membrane interaction. Membrane-active peptides often present facial amphiphilicity, which also promote the interaction with lipid bilayers. To study the role of Trp and facial amphiphilicity in cell interaction and penetration of CPPs, a nonapeptide series containing only Arg, Trp or D-Trp residues at different positions was designed. Our quantitative study indicates that to maintain/increase the uptake efficiency, Arg can be advantageously replaced by Trp in the nonapeptides. The presence of Trp in oligoarginines increases the uptake in cells expressing GAGs at their surface, when it only compensates for the loss of Arg and maintains similar peptide uptake in GAG-deficient cells. In addition, we show that facial amphiphilicity is not required for efficient uptake of these nonapeptides. Thermodynamic analyses point towards a key role of Trp that highly contributes to the binding enthalpy of complexes formation. Density functional theory (DFT) analysis highlights that salt bridge-π interactions play a crucial role for the GAG-dependent entry mechanisms.

## 1. Introduction

Cell-penetrating peptides (CPPs) have attracted much attention in the last two decades for their unique ability to enter cells independently of chirality or receptors. CPPs are short cationic peptides of less than 30 amino acids. There is great diversity in CPP sequences, but they usually contain a high proportion of basic residues, and most particularly arginines [1]. Their uptake mechanism is heavily debated, but it is now accepted that peptide internalization can concomitantly occur by endocytosis and direct translocation through the membrane [2]. These different entry routes suggest that the most abundant molecular species at the plasma membrane, phospholipids and cell surface glycosaminoglycans (GAGs), bind CPPs with different affinity or selectivity. Thus, the analysis of the interactions of CPPs with lipids or GAGs should lead to a better understanding of the molecular mechanisms underlying CPP internalization [3].

Arg residues in CPPs are essential for interactions with negatively charged components of cell membranes. Indeed, on top of electrostatic interactions, guanidinium moieties form bidendate hydrogen bonds with sulfate and carboxylate moieties found on GAGs and phosphates of membrane phospholipid headgroups [4]. Purely cationic CPPs such as polyarginines or Tat could take advantage of these highly favorable interactions and cross the cell membrane through hydrophobic counterion-mediated translocation [4–6]. On the other hand, other basic CPPs such as Penetratin also contain hydrophobic residues, such as Trp residues that are crucial for internalization [7]. In terms of secondary structure, Penetratin is highly versatile since it can be unstructured, adopt α-helical or β-strand conformations depending on its environment (free in solution, bound to membranes, in different cell compartments) and experimental conditions (such as peptide to lipid ratio) [8–13]. It was thus suggested that Penetratin could deeply interact with and perturb the lipid bilayer, and cross the membrane by forming an inverted micelle. Trp residues would help by promoting negative curvature of the bilayer [8]. A simpler analogue of Penetratin, composed of only Arg and Trp was designed [14–16] followed by a shorter 9 residues version to avoid cytotoxicity issues [17]. These R/W peptides are both efficient CPPs, whereas a nine residue R/L peptide is not internalized [18], showing again the essential role of Trp residues.

These designed R/W peptides have the same ability as Penetratin to adopt a facial amphiphilic α-helical secondary structure, as shown both by a peptide secondary structure prediction software [19, 20] and experimental data [18, 21, 22]. The role of facial amphiphilic structuration in CPP internalization has long been debated [23]. For example, the [W48F, W56F]-Penetratin double mutant has stabilized α-helical properties compared to Penetratin [10, 15], while being hardly internalized into cells [7]. A quadruple mutant [I45P, Q50P, M54K, K55P]-Penetratin is still internalized even though it has lost its helical secondary structure [7, 14]. Facial amphilicity has often been used as a key criterion for the design of CPP sequences. For example, it was first believed that the KLA-containing model facial amphiphilic peptide was a CPP due to its amphipathic secondary structure [24, 25]. However, it later appeared that amphipathicity had no direct impact on internalization of the KLA peptide, but could promote binding to intracellular organelle membranes. Strategies exploiting switchable and/or structurally constrained amphipathic helical peptides to control or enhance uptake are still being developed [26–28]. CPPs are often classified according to their ability to adopt a secondary amphipathic structure [23, 29, 30] possibly by analogy with cationic α-helical antimicrobial peptides, another class of membrane-active peptides [31, 32]. For instance, despite its high structural versatility, Penetratin is often classified as an α-helical CPP.

Bechara *et al*. studied the role of Trp and secondary structure in GAG-dependent internalization of various known CPP sequences [33, 34]. In particular, it was found that Trp-containing CPPs tended to adopt β-strand secondary structures and formed large stable aggregates in the presence of GAGs, thus promoting efficient GAG-dependent internalization. In this paper, we aim to push this study one step further by using model nonapeptides containing only Arg and Trp, in order to study the role of the number and the position of the Trp, as well as facial amphiphilicity in the internalization of Arg-rich CPPs.

## 2. Materials and Methods

### Peptide synthesis

All peptides were synthesized using standard Boc solid phase peptide synthesis. Boc-l-Arg(Tos), Boc-l-Trp(For), Boc-Gly, d-Biotin, MBHA Resin (0.53 mmol/g loading) and HBTU were purchased from Iris Biotech GmbH. Boc-(2,2-D_2_)-Gly was obtained from Cambridge Isotope Laboratories. Boc-d-Trp was purchased from Sigma-Aldrich.

d-Biotin was fully oxidized to d-Biotin sulfone (Biot(O_2_), Figure S1) by 4 days treatment with 30% H_2_O_2_ in H_2_O and used without further purification. This avoids further oxidation of the peptide throughout time. Peptides were synthetized by hand on a 0.1 mmol scale for non-deuterated peptides and 0.01 mmol for deuterated peptides. Amino acid (5 eq) activation was performed by HBTU (4.5 eq) in the presence of excess DIEA (12 eq), and Boc deprotection was performed in neat TFA (2× 1 min). Trp side chains were deprotected prior to cleavage by treatment with 10% piperidine in DMF (1, 2, 5, 10, 30 and 60 min successive incubations). Peptides were cleaved from the resin by anhydrous HF (2h, 0°C) in the presence of anisole and dimethylsulfide.

Peptides were purified by reverse phase HPLC on a C18 preparative column (Macherey Nagel) with a H_2_O (0.1% TFA)/MeCN (0.1% TFA) elution gradient. Peptide purity and identity were further characterized by analytical reverse phase HPLC (C18, Higgins Analytical) with a H_2_O (0.1% TFA)/MeCN (0.1% TFA) elution gradient and MALDI-TOF MS (AB Sciex Voyager DE-PRO MALDI TOF or 4700 MALDI TOF/TOF)(Table S1, Figure S2).

### Cell culture, internalization quantification, membrane permeation and cytotoxicity assays

Wild type Chinese Hamster Ovary (CHO-K1, WT, ATCC) and xylose transferase-deficient CHO-pgsA745 (GAG-deficient, ATCC) cells [64] were cultured in Dulbecco’s modified Eagle’s medium (DMEM) supplemented with 10% fetal calf serum (FCS), penicillin (100,000 IU/L), streptomycin (100,000 IU/L), and amphotericin B (1 mg/L) in a humidified atmosphere containing 5% CO_2_ at 37°C.

For internalization quantification assay, 500,000 cells/well are seeded in a 12-well plate the day prior to the experiment so that they reach 1,000,000 on the day of the experiment. The cells are incubated for 1h with 10 µM peptide at 37°C, then extensively washed with Hank’s Buffer Saline solution (HBSS). The cells are then treated with Trypsin, 5 min at 37°C to detach them from the wells and digest the non-internalized and membrane-bound peptide, and trypsin activity is stopped by addition of soybean trypsin inhibitor and bovine serum albumin. The cells are then harvested, the pellets washed and lysed (0.1% Triton X-100, 1M NaCl, 100°C, 15 min) in the presence of a known amount of deuterated peptide acting as an internal standard for MS quantification. The biotinylated peptides are retrieved by incubating the lysate with Dynabeads MyOne Streptavidin C1 (Invitrogen) 1h at room temperature. After several washing steps, the peptides are eluted from the beads by addition of CHCA matrix (saturated in H_2_O:MeCN 1/1, 0.1% TFA). The samples are analyzed in positive ion reflector mode on a Voyager DEPRO MALDI TOF spectrometer (AB Sciex). Quantitty of peptide internalization in CHO cells was quantified by MALDI-TOF MS as extensively previously described [3, 35].

For membrane integrity assay, 5,000 cells are seeded in a 96-well plate the day prior to the experiment, so that they reach 10,000 cells on the day of the experiment. For the cytotoxicity assay, 2,000 cells are seeded so that they reach 4,000 cells on the day of the experiment. The cells are incubated with 10, 25 and 50 µM peptide for membrane integrity and 50 µM peptide for cytotoxicity, for 1h at 37°C. The Cyto-Tox ONE (Promega; LDH release from cells with damaged membranes) and CCK-8 (Dojindo; dehydrogenase activity of viable cells) kits are used according to the manufacturers’ instructions and the plates are read on a FLUOstar microplate reader (BMG Labtech). Briefly, for the CCK-8 kit, after incubation with peptide, the cells are washed with HBSS and 90 µL DMEM + 10 µL CCK-8 solution are added to each well. The absorbance at 450 nm is read after 2h incubation at 37°C. For Cyto-Tox ONE, after incubation with peptide, 100 µL of Cyto-Tox ONE reagent is added to each well. After 10 min at room temperature, 50 µL of stopping solution is added to each well and fluorescence (λ_ex_ = 560 nm, λ_em_ = 590 nm) is immediately measured. 100 % permeation and cytotoxicity was defined with 0.1 % Triton X-100 treatment.

### Sample preparation for calorimetry and ATR-FTIR experiments

Heparin was obtained as a concentrated stock solution (25,000 UI/5mL, 3 mM) from Sanofi and DMPG and POPG from Genzyme as powders.

For liposome preparation, the appropriate amount of lipid is dissolved in CHCl_3_:MeOH 2/1 and a thin film of lipid is formed on the walls of a glass tube by evaporating the solvent with a gentle N_2_ stream. Remaining traces of solvent were further evaporated by leaving the tube 30 min in a desiccator under vacuum. The dried lipids are then resuspended in PBS and vigorously vortexed to form multilamellar vesicles (MLVs). These vesicles can be used as such for DSC experiments. To form large unilamellar vesicles (LUVs) used in ITC experiments, the MLVs are submitted to four freeze-thaw cycles in liquid N_2_/warm water and passed 15 times through a 100 nm polycarbonate membrane using a mini-extruder (Avanti).

For ATR-FTIR experiments, in order to avoid background signal from TFA counterions in peptide, the peptides were dissolved in 1M HCl and freeze-dried. This allowed the exchange of TFA for Cl^−^ counterions.

### Calorimetry assays

All ITC experiments were performed with a NanoITC calorimeter (TA instruments) and data were analyzed using NanoAnalyze (TA instruments) using a simple binding model with *n* independent binding sites. The volume of the calorimetric cell is 983 µL and the injection syringe is 250 µL.

Isothermal Titration Calorimetry (ITC) experiments were performed on a NanoITC calorimeter (TA Instruments). 250 µL of a Heparin solution (30-75 µM in PBS) was injected by 1×2 µL and 24×10 µL steps into 1 mL peptide solution (30-75 µM in PBS). For lipid binding experiments, 250 µL of a 1-palmitoyl-2-oleoyl-*sn*-glycero-3-phospho-(1’-rac**-**glycerol) (POPG) large unilamellar vesicle (LUV) suspension (100 nm, 2-5 mM lipid concentration in PBS) was injected by 1×2 µL and 24×10 µL steps into 1 mL peptide solution (50-100 µM in PBS). Binding parameters were determined by using the one site binding model provided by the NanoAnalyze software (TA Instruments). Representative ITC data injections of heparin (HI) or POPG LUVs into peptide solutions are shown on figure S4. Experiments were performed twice and the numbers given in tables 2 and 5 of the manuscript are averaged on the two experiments.

**Table 1:**
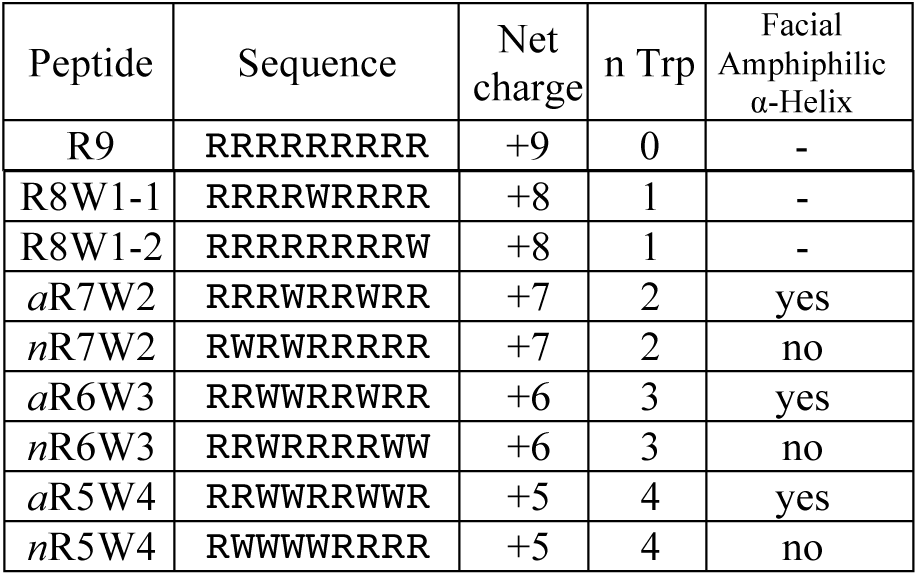
Peptide sequences used in the study. All peptides are amidated at the *C*-terminus and carry a Biot(O_2_)-GGGG tag at their *N*-terminus.

**Table 2:** Peptide binding to heparin studied by ITC.

| Peptide | $K_D$<br>nM | $\Delta H$<br>kcal/<br>mol | $T\Delta S$<br>kcal/mol | $n$<br>peptide/<br>heparin | % of<br>charge<br>equilib-<br>ration |
| --- | --- | --- | --- | --- | --- |
| R9 | 7 | -71.7 | -60.5 | 8 | 72 |
| R8W1-1 | 30 | -81.7 | -71.5 | 10 | 80 |
| R8W1-2 | 21 | -92.3 | -82.0 | 7 | 56 |
| aR7W2 | 18 | -96.8 | -86.3 | 11 | 77 |
| nR7W2 | 26 | -93.5 | -82.9 | 12 | 84 |
| aR6W3 | 15 | -116.4 | -105.6 | 9 | 54 |
| nR6W3 | 23 | -123.3 | -113.0 | 11 | 66 |
| aR5W4 | 25 | -176.6 | -165.9 | 17 | 85 |
| nR5W4 | 25 | -179.3 | -169.0 | 14 | 70 |

**Table 3:** Peptide binding to POPG LUVs studied by ITC.

| Peptide | $K_D$<br>($\mu\text{M}$) | $\Delta H$<br>(kcal/mol) | $T\Delta S$<br>(kcal/mol) | $n$ (lipid /<br>peptide) |
| --- | --- | --- | --- | --- |
| *R9* | 1.7 | -0.60 | 7.4 | 8.2 |
| *R8W1-1* | 7.8 | -0.72 | 6.2 | 7.0 |
| *R8W1-2* | 3.2 | -0.79 | 6.7 | 6.5 |
| *aR7W2* | 6.4 | -0.84 | 6.2 | 5.6 |
| *nR7W2* | 3.4 | -0.81 | 6.7 | 7.2 |
| *aR6W3* | 6.8 | -1.39 | 5.7 | 5.2 |
| *nR6W3* | 2.3 | -1.27 | 6.5 | 5.6 |
| *aR5W4* | 2.9 | -2.53 | 5.0 | 3.5 |
| *nR5W4* | 2.3 | -2.10 | 5.7 | 3.2 |

**Table 4:** Secondary structure (ATR-FTIR) of R5W4 peptides in the absence or presence of heparin (8.3 µM heparin and peptide concentration according to the stoichiometries determined by ITC).

| Secondary structure element | Wave-numbers (cm<sup>-1</sup>) | Percentage of structural elements |  |  |  |
| --- | --- | --- | --- | --- | --- |
|  |  | aR5W4 |  | nR5W4 |  |
|  |  | Alone | +Hep | Alone | +Hep |
| β-strand | 1627 | 11 | 20 | 17 | 21 |
| Random coil | 1643 | 24 | 20 | 22 | 22 |
| α-Helix | 1659 | 29 | 30 | 26 | 26 |
| Turn | 1679 | 36 | 30 | 35 | 32 |

**Table 5:** Binding of R6DW3 to Heparin and POPG.

| | $K_D$ | $\Delta H$<br>(kcal/mol) | $T\Delta S$<br>(kcal/mol) | Stoichiometry |
| --- | --- | --- | --- | --- |
| R6DW3/<br>HI | 16.5 nM | -91.1 | -80.5 | 11<br>peptide/heparin |
| R6DW3<br>/POPG | 9.5 μM | -0.74 | 6.2 | 6.4<br>lipid/peptide |

Differential Scanning Calorimetry (DSC) experiments were performed on a NanoDSC calorimeter (TA Instruments). To a suspension of 1,2-dimyristoyl *sn*-glycero-3-phospho-(1’-rac**-**glycerol) (DMPG) multilamellar vesicles (MLVs) (1 mg/mL lipid concentration; 1.47 mM in PBS), peptide was added to reach a peptide-to-lipid molar ratio of 1/50. Samples were scanned at 1°C/min between 0°C and 50°C at least 3 series of alternated heating and cooling scans.

### Peptide secondary structure studied by IR spectroscopy

Attenuated Total Reflection (ATR) FTIR spectra were recorded on a Nicolet 6700 FT-IR spectrometer equipped with a liquid nitrogen cooled mercury-cadmium-telluride detector (ThermoFisher Scientific), with a spectral resolution of 4 cm^−1^. Two hundred interferograms, representing were co-added. The peptides were deposited on the Ge crystal at 100 µM in PBS buffer. The peptide/heparin mixtures were prepared using the stoichiometries determined by ITC. To determine the secondary structure element of each peptide or protein, spectra were analyzed with an algorithm based on a second-derivative function and a self-deconvolution procedure (GRAMS and OMNIC softwares, ThermoFisher Scientific) to determine the number and wavenumber of individual bands within the spectral range 1720-1500 cm^−1^. Example of Amide I band deconvolution for *a*R5W4 peptide is shown on figure S6 and secondary structures extracted from ATR spectra for all peptides are given in Table S2.

### DFT analyses

The energies of all complexes included in this study were computed at the BP86-D3/def2-TZVP level of theory. The calculations were performed with the TURBOMOLE version 7.0 program [36]. No constrains have been imposed during the optimizations. The minimum nature of the complexes has been confirmed by performing frequency calculations at the same level. For the calculations we used the BP86 functional with the latest available correction for dispersion (D3) [37]. The MEP (Molecular Electrostatic Potential) surfaces have been computed at the B3LYP/6-31+G* level of theory by means of the SPARTAN software [38]. Values have been plotted onto the van der Waals isosurface (0.001 a.u.) unless otherwise noted. In order to reproduce solvent effects, we have used the conductor-like screening model COSMO [39], which is a variant of the dielectric continuum solvation models [40]. We have used water as solvent.

## 3. Results

### 3.1. Peptide design

We designed a series of 9 nonapeptides composed of Arg and Trp exclusively. We varied the number of Trp from 0 to 4 and introduced them at different positions in the sequence, so that the peptides can potentially adopt an amphipathic α-helical secondary structure, or not. All the peptide sequences are presented in Table 1. Each peptide is referred to by its number of Arg and Trp, preceded by *a* for peptides with a potential facial amphiphilic structure and *n* for peptides without potential facial amphiphilic structure. Predictions for facial amphiphilic structures were obtained with the software PEP-FOLD3 [19, 20]. For example, *a*R7W2 refers to a peptide with 7 Arg and 2 Trp and potential for a facial amphiphilic structure, *n*R7W2 to a peptide with 7 Arg and 2 Trp and no potential for a facial amphiphilic structure (Fig. 1; See SI for the other peptides). For peptides with only one Trp, as facial amphiphilicity is not clearly defined, peptides were distinguished by numbers (R8W1-1 and R8W1-2, see Table 1). *a*R6W3 corresponds to a peptide that we have already extensively studied and previously referred to as (W/R) [17], (R/W)9 [2], RW9 (18], R_6_W_3_ [33], R6/W3 [41]. All peptides are amidated at the *C*-terminus and bear a Biot[O_2_)-GGGG tag at their *N*-terminus for intracellular quantification purposes [35].

**Figure 1:**
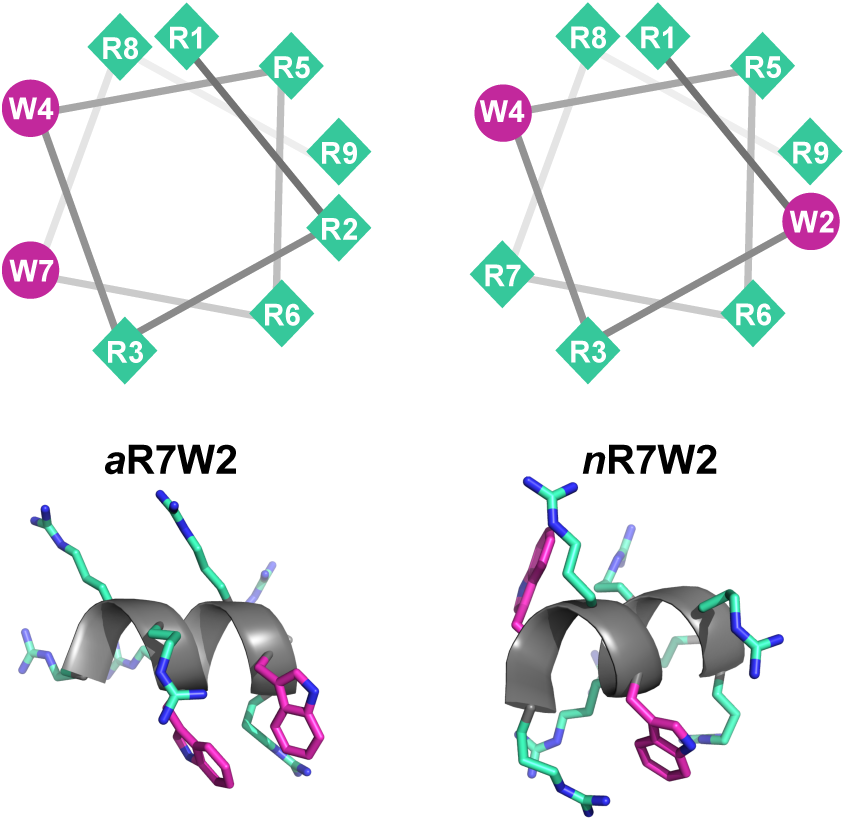
Helical wheel projections and simulated secondary structures using the structure prediction software PEP-FOLD3 for *a*R7W2 and *n*R7W2 peptides, showing facial amphiphilicity (*a*) and non-facial amphiphilicity (*n*). See SI for the other peptides.

### 3.2. Peptide internalization

We studied the internalization of all nine peptides in CHO-K1 (WT) and CHO-pgsA 745 (deficient in chondroitin (CS) and heparan sulfates (HS) GAGs) cells at 37 °C.

#### 3.2.1 Peptide internalization in WT cells

The amounts of internalized peptide in WT cells are presented in Fig. 2A. When comparing two peptides with the same number of Trp but at different positions in the sequence, it appears that Trp position has no impact on the amount of internalized peptide, except in the case of the peptides with 4 Trp. In this case, facial amphiphilicity strongly favors peptide uptake (Fig. 2A). When comparing all facial amphiphilic peptides (Fig. 2B), the number of Trp has a great influence on internalization efficacy. In particular, *a*R5W4 is 3-fold more efficiently internalized than *a*R6W3. On the other hand *a*R7W2 is significantly less internalized than *a*R6W3 and *a*R8W1. Finally, there is no significant difference between *a*R8W1 and R9. When comparing all non-facial amphiphilic peptides (Fig. 2C), the number of Trp also has a strong influence, though there is no significant difference between *n*R5W4 and *n*R6W3. In this series, *n*R7W2 is also the least efficient.

**Figure 2:**
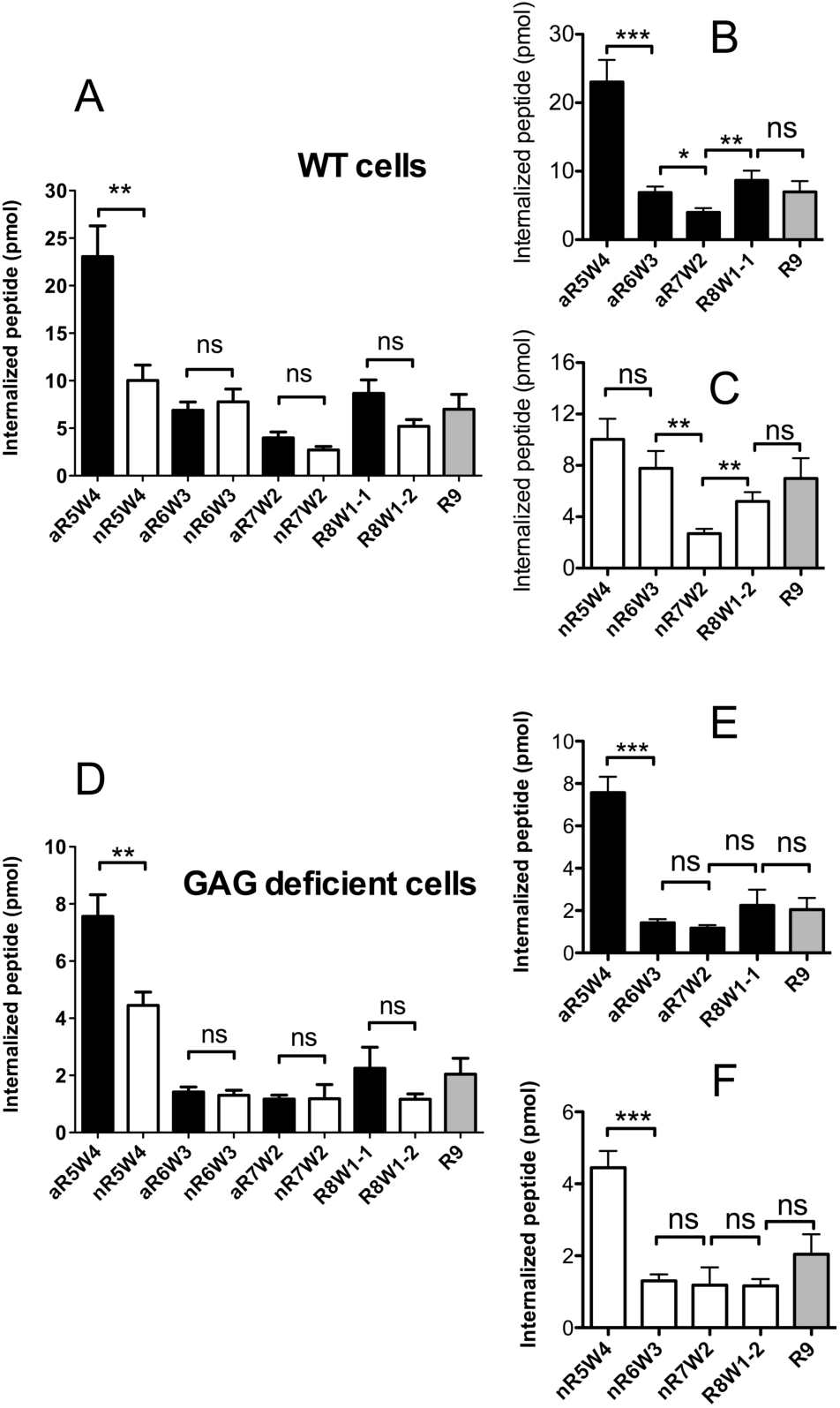
Peptide internalization after incubation of 10 μM (1 mL) peptides at 37°C with 10^6^ WT (A, B, C) or GAG deficient cells (D, E, F) for 1 hr. Panels A and D show the quantification for all peptides (black: facial amphiphilicity (*a*), white: non-facial amphiphilicity (*n*)). Panels B and E focus on peptides with facial amphiphilicity (*a*), panels C and F on peptides with non-facial amphiphilicity (*n*). Significance was tested using Welch’s t-test comparison of two columns (ns p>0.05, * 0.05>p>0.01, ** 0.01>p>0.001, *** p<0.0001). Each experiment was repeated at least three times independently and in triplicates. Error bars represent standard error (SEM).

#### 3.2.2 Peptide internalization in GAG deficient cells

The amount of internalized peptides in CS and HS GAG deficient cells are presented in Fig. 2D. Overall, the amounts of intact peptide detected in cells is 3 to 5-fold lower than in WT cells, showing that HS and CS-type GAGs are involved (though not strictly necessary) in the internalization process for these peptides. As observed for WT cells, facial amphiphilicity only impacts the uptake of peptides containing 4 Trp. In terms of number of Trp in the sequence, it appears that GAG-independent peptide internalization is less sensitive to this number compared to entry in WT cells. In particular, no significant difference between 3, 2, 1 or 0 Trp could be observed, whatever the peptide series (*a* or *n*) (Fig. 2E, 2F).

### 3.3 Peptide-induced cell-membrane permeabilization and cytotoxicity

We also checked whether the peptides were cytotoxic or induced membrane permeabilization on WT cells (Fig. 3). Overall, all peptides were not or little permeabilizing at 10 and 25 µM (less than 10% leakage). At 50 µM, most peptides still had little permeabilizing effects (less than 15%), with the notable exception of R5W4 inducing 30% leakage. We also tested the peptides cytotoxicity at 50 µM (Fig. 3). It closely matched the membrane integrity data. It is worth highlighting that the peptide containing 4 Trp, which compromises membrane integrity is also the one with the higher internalization rate. However, in the internalization assay the peptide to cell ratio, P/C=10-14 that corresponds to 10 µM peptide incubated with106 cells in 1 mL, is 10-100 times lower than in the case of cytotoxicity and permeabilization assays (P/C between 2.5×10^−13^−10^−12^ for cytotoxicity: 10-50 µM peptide incubated with 10^4^ cells in 100 µL; P/C between 10^−13^-5×10^−13^ for permeabilization: 10-50 µM peptide incubated with 4,000 cells in 100 µL). It suggests that although used at a non cytotoxic concentration, the significant increase in internalization observed for R5W4 peptide in GAG-deficient cells, could arise from slight membrane permeabilization, such as transient pore formation. Altogether, it appears that 3 Trp is a maximum number for efficient internalization of these nonapeptides in cells without associated cytotoxicity at the concentrations used for internalization.

**Figure 3:**
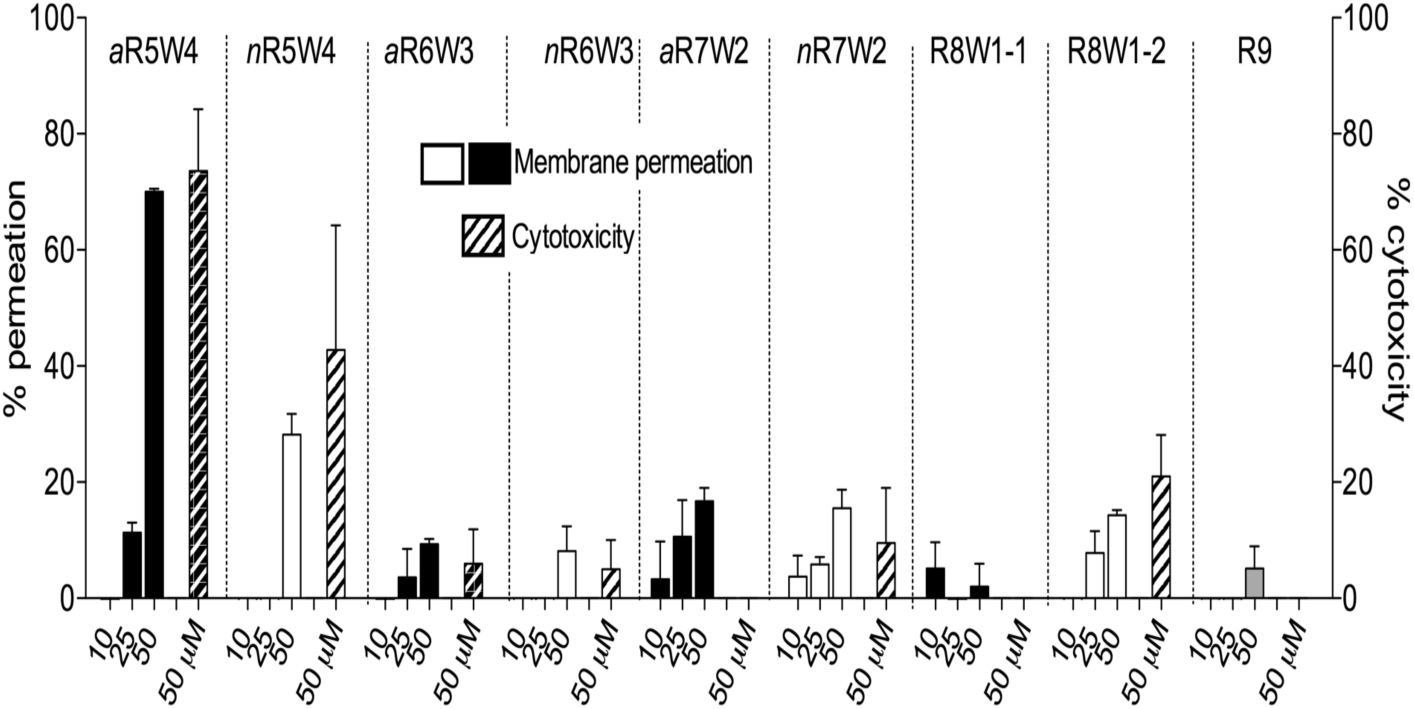
Membrane leakage upon incubation for 1 hr at 37°C with 10 µM, 25 µM and 50 µM (100 μL) peptide (black facial amphiphilic, white non-facial amphiphilic) and cytotoxicity with 50 µM (100 μL) peptide (hashed) assayed on respectively 10,000 and 4,000 WT cells. Experiments were repeated independently at least two times in triplicates. Error bars represent standard error (SEM).

### 3.4 Peptide binding to GAGs

As GAGs are obviously important partners for CPP internalization, we studied the direct interaction between the nine peptides and heparin (taken as a GAG mimic) by ITC. Results are displayed in Table 2. K_D_ values were all in nM range, with no obvious difference between sequences. Binding to heparin was always associated with large favorable enthalpies, the absolute value increasing with the number of Trp in the sequence and decreasing with the number of positive charges. This shows that peptide/GAG binding is far from being purely electrostatic. At the same time, peptide binding to heparin is associated with unfavorable entropies, possibly due to the restriction of the number of accessible conformations of heparin chains upon peptide binding, as previously reported for other Trp-rich CPPs [33]. Finally, the number of peptides bound per heparin chain increases with the number of Trp in the sequence, and corresponds to 54 to 85% of charge equilibration, considering an average of 100 negative charges per heparin chain. No obvious correlation between amounts of internalized peptide and thermodynamic parameters of binding to heparin could be derived.

### 3.5 Peptide binding to lipids

We then studied the binding of the peptides to POPG large unilamellar vesicles. We chose POPG, even though this lipid is not found in the outer leaflet of eukaryotic cells. Previous studies have shown that CPPs bind only loosely to PC membranes [42] and that negative charges are required to trigger measurable peptide/membrane interactions [18]. We thus chose POPG as a simple mimic of the negatively charged lipids that could be found in the membrane. Results are displayed in Table 3. All peptides bound to POPG with an apparent K_D_ in the low µM range. Peptide binding to POPG was enthalpically and entropically favorable, with a major entropic contribution, as already previously reported for *a*R6W3 [43]. The large positive entropy of binding could result from the increase in lipid disorder, counterions release from the charged POPG and peptide, or the decreased amount of polarized water between the two charged molecules, most probably a combination of all. Binding enthalpy increases with the number of Trp in the sequence, as was previously reported for other Trp-containing CPPs [44]. The stoichiometry of binding is close to simple charge equilibration, provided the peptides can translocate through the POPG bilayer. This is the case for *a*R6W3 and R9 for instance [45, 46]. However, only a fraction of the CPP enters into lipid vesicles, and it is more likely that the observed stoichiometries are mainly due to incomplete ion pairing on the outer leaflet of the vesicle, as was already previously suggested for Arg-rich CPPs [4, 5, 46, 47]. In addition, the number of lipids per peptide linearly decreases according to the reduction of positive charges (or the increase in the number of Trp). As for GAG binding, no obvious correlation between internalization and lipid binding could be observed. Also, it appears that the position of the Trp in the sequences has little effect on the binding parameters.

### 3.6 Peptide effect on bilayer organization

To further investigate the interaction of the peptides with lipid membranes, we studied the effect of the peptide on the phase transitions of DMPG. Results obtained for a peptide-to-lipid ratio of 1/50 are shown on Fig 4.

**Figure 4:**
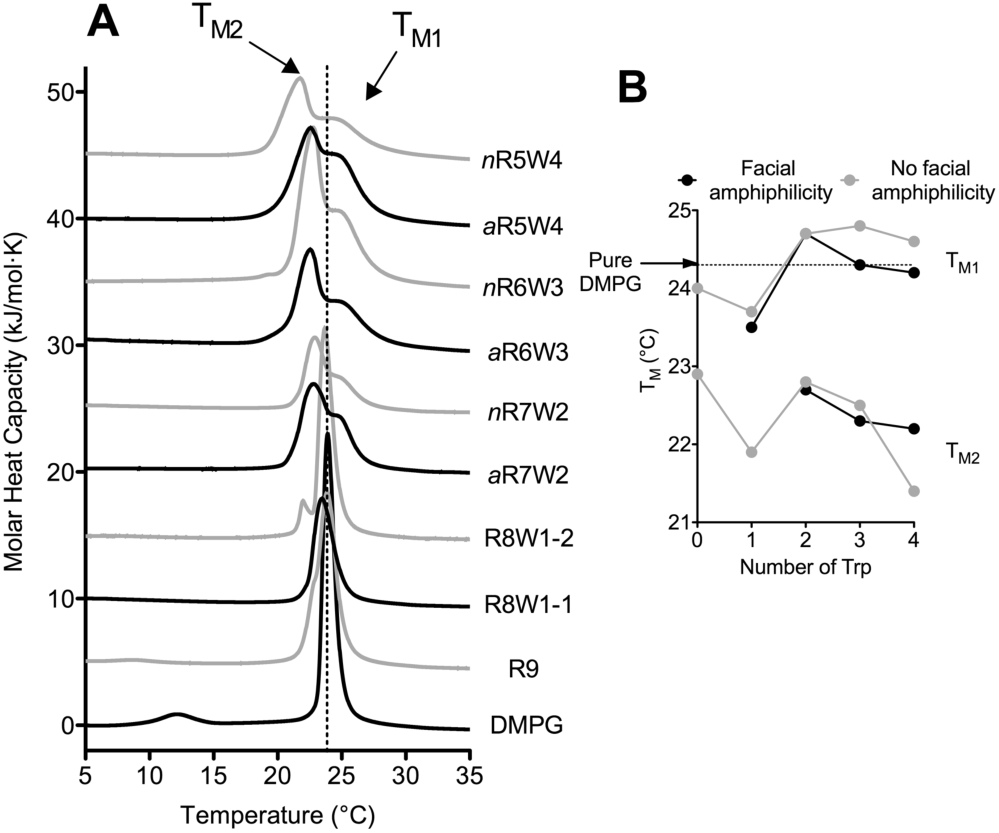
Perturbation of membrane organization by CPP addition probed by DSC. (A) Thermograms showing the phase behavior of DMPG MLVs in the presence of peptides at P/L = 1/50. (B) Phase transition temperature(s) according to the number of Trp in the peptide sequences. The dotted lines mark the main phase transition temperature of pure DMPG. Thermograms were obtained from 5 heating / cooling cycles.

Overall, not surprisingly, addition of peptide leads to the disappearance of the pre-transition, suggesting an interaction between the peptides and the lipid headgroups, as already reported for many Arg-rich CPPs [18, 48]. As regards to the main transition, in most cases, a splitting of the peak can be observed. This suggests lipid phase separation, with more fluid low T_M2_ regions, strongly affected by the presence of the peptide, and higher T_M1_ regions, closer or slightly higher than that of the lipid alone. Globally, the value of the low T_M2_ decreases with the number of Trp in the sequences showing that increasing the number of Trp will lead to more perturbation of the peptide-enriched regions. The borders of these segregated domains of different fluidity are suggested to be regions of enhanced membrane permeability [49] and are proposed to be an entry route for CPPs through the plasma membrane [34, 50].

### 3.7 Peptide secondary structure and secondary structure disruption

#### 3.7.1 Secondary structure determination by polarized ATR-FTIR

The peptides in our series were designed so that some can adopt a facial amphiphilic α-helical structure in the presence of lipids and/or GAGs, and some cannot. We studied the secondary structure of these peptides by ATR-FTIR in the presence or absence of heparin. Globally, these peptides showed great structural plasticity, suggesting they can structurally adapt to different types of binding partners. Example for *a*R5W4 and *n*R5W4 are given in Table 4. Other data are provided in SI. Altogether, this structural plasticity cannot be directly related to the internalization efficiency of the peptides. In particular, although the peptides can all adopt a α-helical structure, we could not conclude whether it is mandatory for internalization.

#### 3.7.2 Secondary structure disruption

Since the peptides display great structural versatility, one cannot predict whether α-helical structures could be selected for or during the internalization steps. In order to further assess the role of facial amphiphilicity, and more generally of secondary α-helical structure in CPP internalization, we synthesized the peptide R6DW3 (RRDWDWRRDWRR) that combines L-Arg and D-Trp, and tested its internalization in WT cells and interactions with potential membrane partners, as described above for the other nonapeptides. Indeed, introducing D-amino acids in the middle of a peptide sequence prevents α-helix formation [51].

Strikingly, R6DW3 was internalized to the same extent as its all-L amino acids analogue *a*R6W3 in WT cells, confirming that a well-defined secondary structure, in particular one displaying facial amphiphilicity, is not an important feature for CPP internalization (Fig. 5A). Binding parameters of R6DW3 to heparin and POPG as determined by ITC also fall within the same range as *a*R6W3 (Table 5). The effect of R6DW3 on a DMPG bilayer organization, as probed by DSC is almost identical to that of *a*R6W3 (Fig. 5B). These results show clearly that facial amphiphilicity is not required for peptide binding and insertion into membranes, and further efficient internalization.

**Figure 5:**
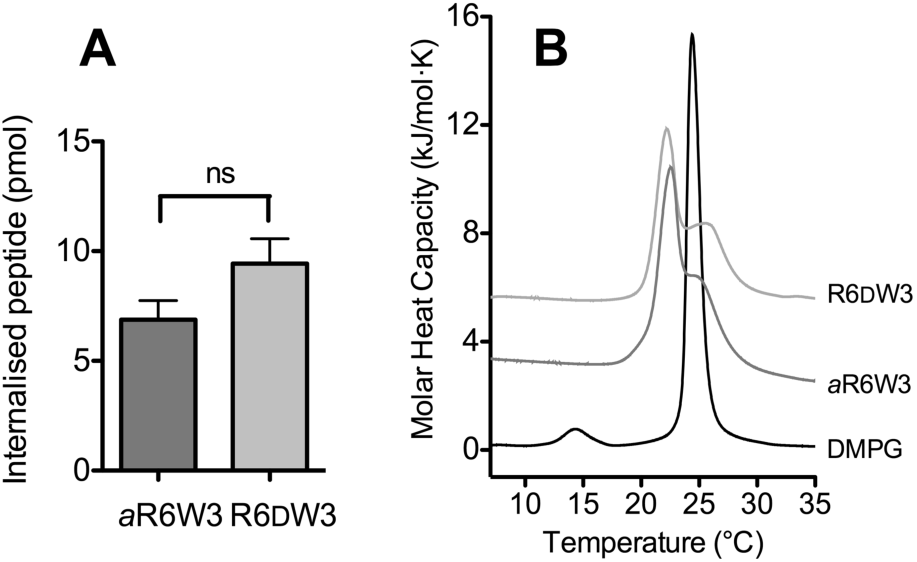
Compared internalization rate in WT cells (A) and effect on a DMPG membrane probed by DSC (B) of *a*R6W3 and R6DW3.

### 3.8 DFT modeling

#### 3.8.1 Anion-π and salt-bridge π modeling

To further understand the role of Trp in the interaction with sulfated polysaccharides, in particular in the highly favorable enthalpy values we measured, we have next computed the complexes of 3-methyl indole as a model of the side chain of tryptophan, with a cation (formamidinium), anion (formate) and also a salt bridge formamidinium–formate. The results are given in Fig. 6 where it can be observed that the formation of the cation-π complex is favorable (Fig. 6C, –5.2 kcal/mol). However, the optimization starting from the anion-π complex leads to the formation of a H-bonded complex (Fig. 6B, –5.4 kcal/mol), as expected. If the anion-π assembly is imposed (by fixing the geometry), the resulting interaction energy is unfavorable (Fig. 6A, +8.4 kcal/mol). In contrast, the complex with the salt-bridge formamidinium–formate is favorable in –4.8 kcal/mol (Fig. 6D). This preliminary study anticipates that the role of Trp in the binding to GAGs is not due to anion–π interactions. We have then further analyzed the interaction of model Arg/Trp or Arg/Gly tri- and tetrapeptides with GAG motif disaccharides.

**Figure 6:**
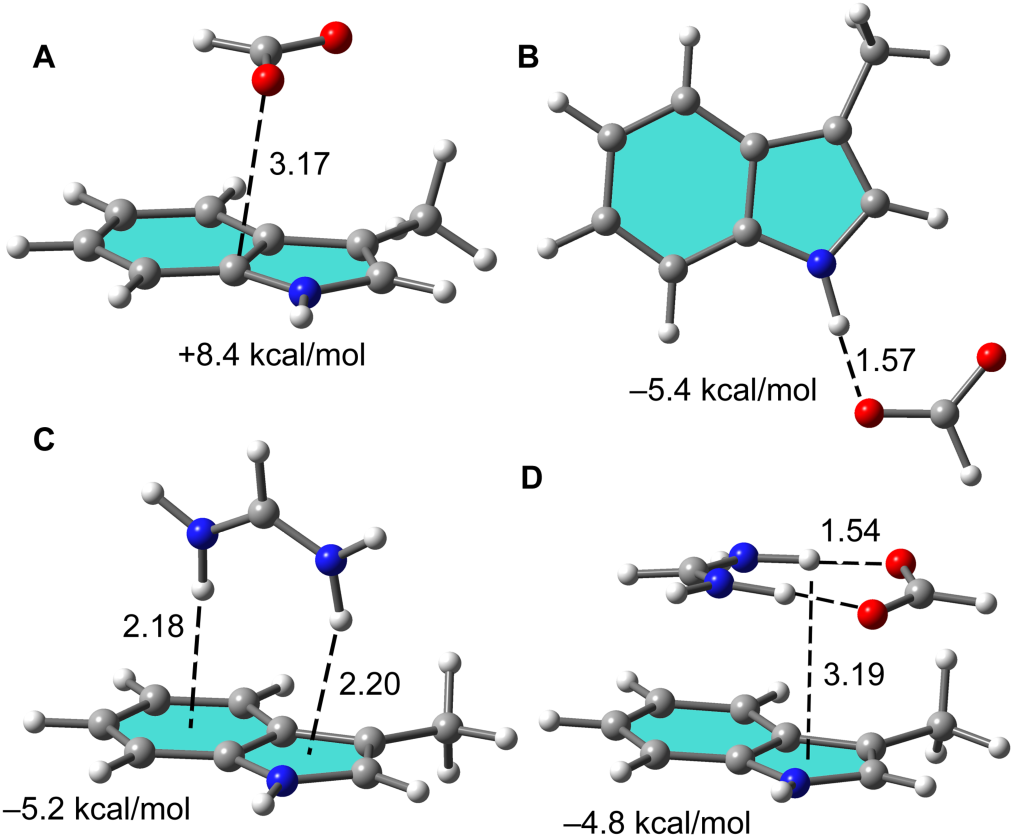
Optimized geometries of 3-methylindole with formate anion (A, B), formamidinium (C) and the salt-bridge (D). Distances in Å. The interaction energies are also indicated.

#### 3.8.2 Interactions with GAG models

In Fig. 7A we show the minimalistic models of the CPPs and GAGs used for the DFT analysis. We have used the monosulfated disaccharide D1 as a model of chondroitin 4-sulfate (CS-A) GAGs. We have methylated the bridging O atoms. We have used both RWWR and RWR motifs as models for the CPPs. In addition, to investigate the role of Trp in the binding mechanism, we have replaced Trp by Gly in RGGR and RGR peptides.

**Figure 7:**
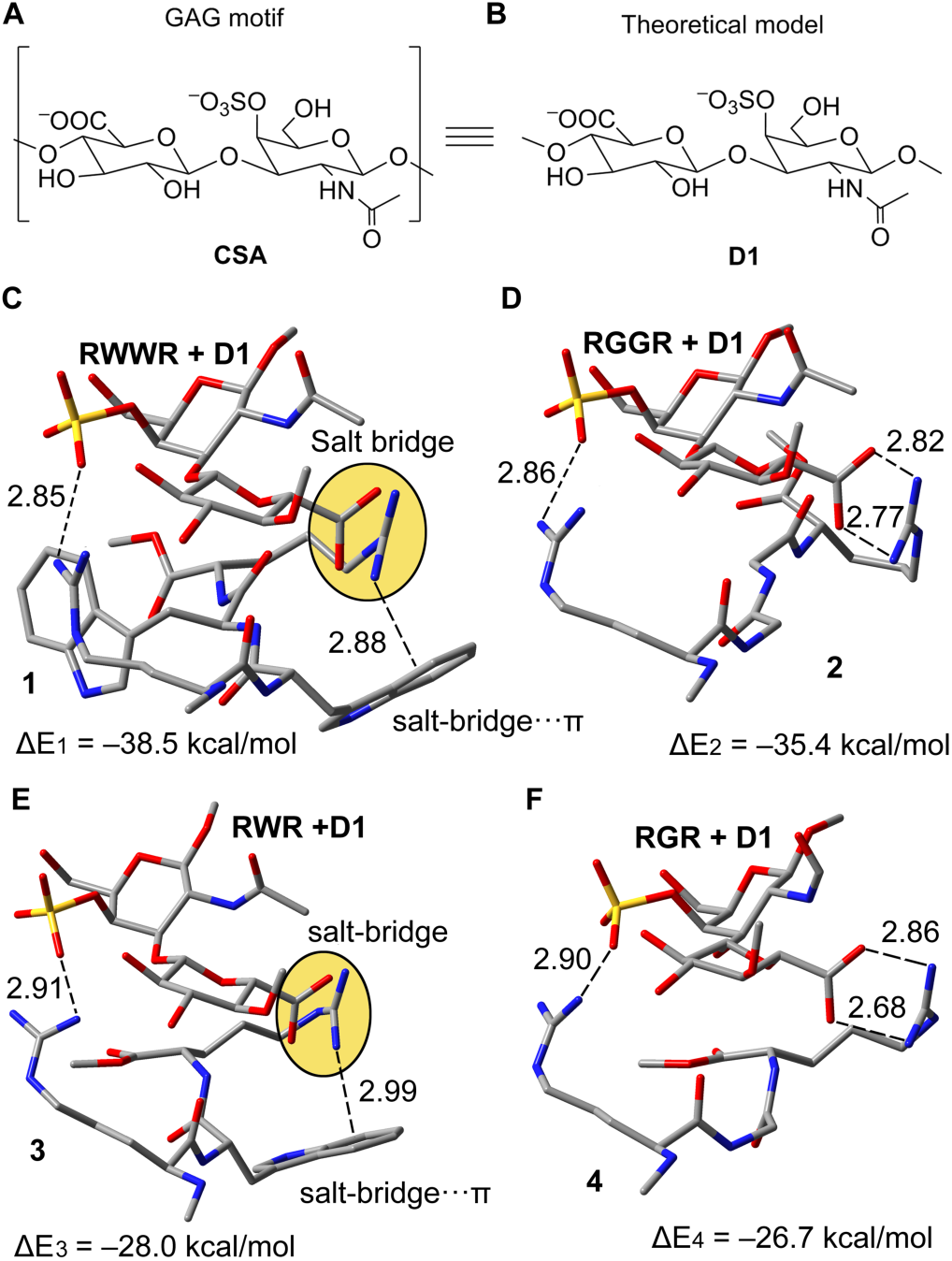
(A, B) GAG model used for simulations. (C, D) Optimized complexes **1** (RWWR+D1) and **2** (RGGR+D1). (E, F) Optimized complexes **3** (RWR+D1) and **4** (RGR+D1). Distances in Å. Salt-bridge distance measured from the N atom to the ring centroid of the six membered ring. H-atoms omitted for the sake of clarity.

The geometries and binding energies for the D1 series are shown in Fig. 7. The analysis reveals the presence of salt bridge interactions in both the RWWR and RWR complexes with D1. Remarkably, the salt bridge that is established between the carboxylate group of the disaccharide and the side chain of R is located over the Trp side chain π-system, thus generating a salt-bridge-π interaction. In the analog where Trp has been replaced by Gly, the interaction energy is reduced in 3 kcal/mol for the RGGR and in 1.3 kcal/mol in RGR, thus supporting the importance of the indole group in the Trp side chain in the affinity of RWWR and RWR sequences to D1. It is also interesting to note that both positive guanidinium groups interact with the negative groups of the disaccharide thus explaining the large interaction energies. It is also worth mentioning that the RWWR sequence has more affinity to D1 than the RWR one in almost 10 kcal/mol.

#### 3.8.3 Electrostatic potential surfaces

Finally, we computed the electrostatic potential surfaces for 3-methyl-indole and its complex with the salt-bridge, which are shown in Fig. 8. It can be observed that the presence of the ion-pair significantly affects the polarization of the π-system. In fact, the MEP values are negative over both six and five membered rings of the 3-methylindole moiety. Upon complexation of the ion-pair, the electronic charge distribution of the system significantly changes creating an induced dipole. The view from above in Fig. 8C, where only the π-surface is represented clearly reveals that the otherwise reddish color of the MEP surface in the free π-system (Fig. 8A) changes to green over the cation (slightly positive) and orange over the anion (slightly negative), thus revealing a significant polarization of the indole moiety. In fact, the dipole moment of the unperturbed chromophore increases from μ = 2.0 D to 4.9 D upon complexation.

**Figure 8:**
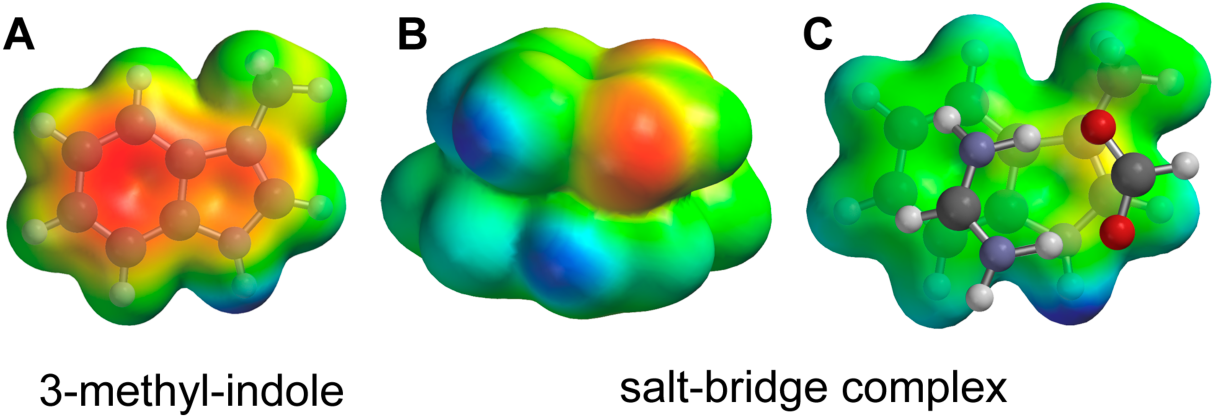
(A, B) Electrostatic potential surfaces of 3-methyl-indole and the complex (Isovalue 0.001 a.u.). (C) Electrostatic potential surface of the π-surface only. Red= negative, blue= positive, isovalue 0.09 au/± 50 kcal/mol.

## 4. Discussion

Arg-rich CPPs have been studied for a long time by different teams. Internalization mechanisms of Arg-rich CPPs remain a puzzle though the new combination of cell studies, biophysics and molecular simulation yield very interesting insight on possible original translocation mechanisms [52]. In the present study, we analyzed the effect of the number of Trp by replacing Arg residues in nonapeptides. On the one hand, Mitchell and Wender previously reported that decreasing the number of Arg from 9 to 5 in oligoarginine sequences led to a stepwise and linear decrease in internalization efficiency of the corresponding peptides [53, 54]. On the other hand, Rydberg *et al*. also investigated the number and position of Trp residues in polyarginine sequences, keeping the number of Arg residues constant (eight) and increasing the number of Trp stepwise [21, 55]. Herein we kept the peptide length to 9 residues and replaced gradually Arg with Trp. As shown in Fig. 9, Trp can compensate for the loss of positive charges very efficiently: R6W3 peptides have similar internalization efficacy as R9, while R6 is only 20% active compared to R9 [53, 54]. This result is consistent with the work of Mishra *et al*. who previously reported that addition of a single aromatic group (Trp, Fluorescein) drastically impacts the translocation mechanism of R6 peptide [56]. Although a compensation for Arg loss by Trp addition is observed in both WT and GAG-deficient cells, in WT cells peptide internalization was actually potentiated by the presence of Trp, in the absence of cytotoxicity.

**Figure 9:**
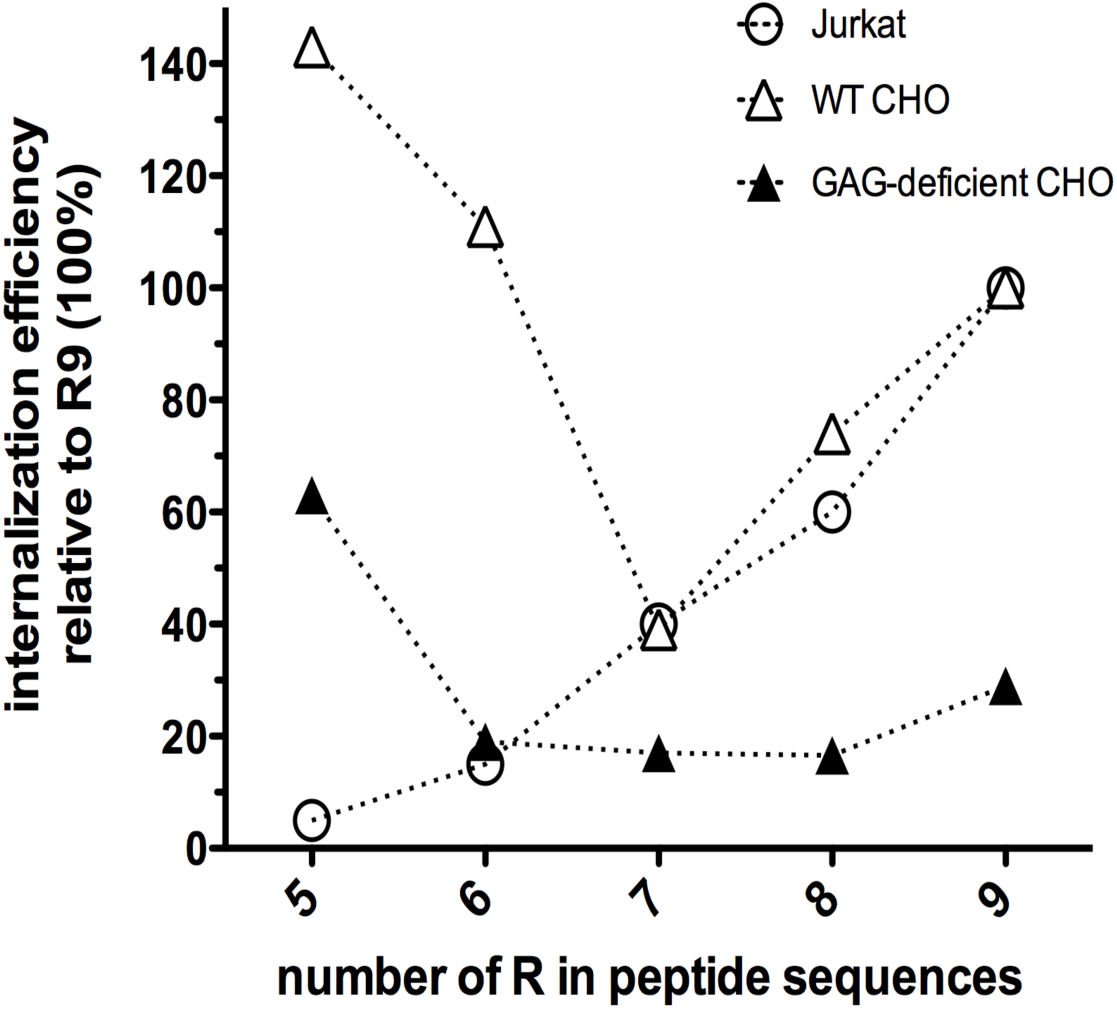
Internalization efficiency of oligoarginines Rx (white circles) and *n*RxWy (black triangles, white triangles) peptides normalized to R9, where x+y = 9. Data for Rx peptides were taken from (53, 54).

Globally, our findings are consistent with the results obtained by Rydberg *et al.* [21, 55], that is increasing the number of Trp leads to better uptake together with an increased cytotoxicity and membrane permeation, with a threshold at 4 Trp. Rydberg *et al.* also show that the affinity of their peptides for membrane models all fall within the same range, making this parameter a bad predictor for cell uptake. However, they found that the position of the Trp in the sequence is an important factor, which is apparently in contradiction with our findings. Interestingly, we do see an effect of the Trp position, but only with the R5W4 peptides, when Rydberg *et al.* only studied the role of Trp position for the sequences containing four Trp. Their study was mainly focused on backbone spacing whereas we mainly focused on potential facial amphiphilicity. Their most efficient sequence is the peptide RWmix (RWRRWRRWRRWR) that could potentially adopt a secondary structure with the Trp facing on the same side of an helix (though with uncomplete facial amphiphilicity), but it remains unclear whether backbone spacing or facial amphiphilicity is the important factor. Taken together, our findings and those described by Rydberg *et al*. both point towards a key role of Trp in Arg-rich CPPs cell uptake, four Trp appearing as a breaking point leading to substantially increased uptake and toxicity, at least for peptide length from 9 to 12 residues. For peptides with three Trp or less, our data clearly prove that facial amphiphilicity is not a prerequisite for efficient internalization as it is often claimed [11]. We also show that facial amphiphilicity has little impact on the binding of R/W CPPs to GAGs or lipids, and on the way they affect the organization of a lipid bilayer. Quite the opposite, it seems that for these CPPs an undefined, plastic and adaptable structure could be an asset for cell penetration, both for GAG-dependent and independent pathways. This structural plasticity of peptides has recently been highlighted as a crucial parameter to discriminate cell-penetration properties through non-endosomal mechanisms from cell toxicity [57]. Such plasticity could explain the apparent discrepancy reported by two independent works on penetratin structure in cells observed by Raman microscopy. In the study of Ye *et al.*, the authors report that Penetratin is random coil or adopts a β-strand structure in the cytoplasm and a β-sheet structure in nucleus of metastatic melanoma SK-MEL-2 cells [12]. In the study of Fleissner *et al.*, using myoblast C2C12 cell line, the authors unveiled a α-helical conformation of penetratin in the cytoplasm and a β-sheet structure in the nucleus [13]. This apparent discrepancy could arise from the difference in cell-surface and organelle membrane partners in the two cell lines.

Counter-intuitively, the number of Trp in our sequences essentially impacts cell uptake in WT cells (*ie* mostly GAG-dependent uptake), whereas it has less impact on the uptake in GAG deficient cells, where the membrane lipids are likely more accessible. Trp is generally considered as a hydrophobic residue that would be expected to promote peptide/lipid interactions. Increasing the number of Trp in the sequences indeed leads to more favorable binding enthalpies when investigating CPP/lipid interactions. In addition, in membrane proteins Trp is found in the electrostatically complex lipid bilayer/water interface, a location likely resulting from its unique π electronic structure and quadrupolar moment. Yet, increasing the number of Trp leads to even more favorable binding enthalpies when studying CPP/heparin interactions. Previous work from our group had already pointed towards an important role of Trp in CPP/GAG interactions that we hypothetically attributed to potential π-anion interactions [33]. The improvement of Trp in the binding to GAGs of cationic peptides has also been observed previously for other types of peptides, but the authors did not give any explanation for such binding impact [58]. Interestingly, targeting GAGs has been recently demonstrated as an efficient strategy for cell delivery purposes [59].

Recently, Matile’s group elegantly reported small anionic amphiphiles, such as pyrene butyrate, as potent oligoarginines binders through ionpair-π interactions, resulting in activation of the cell-penetration for these peptides [60, 61]. Herein we go one-step further by showing that ionpair-π interactions can occur between Arg/Trp-containing peptides and carboxylate groups in GAGs, due to the unique polarizability of Trp, that is much higher than for any other aromatic amino acid (Phe, Tyr or His). These ionpair-π interactions occurring in biological systems could potentiate CPP internalization. Our simulation studies show that this type of interactions can definitely occur between RWWR/RWR motifs and GAGs. The higher energy for the RWWR vs. RWR motif is totally consistent with the experimental ITC data. Whereas Chuard *et al*. evidenced the role of ionpair-π in CPP internalization using synthetic aromatic systems with tuned electronic properties (push-pull dipoles) [60], we show the same type of mechanism is occurring in natural peptide sequences, using the unique properties of Trp, the most electron-rich and polar natural aromatic amino-acid.

From the present and other past studies from our group and others, we finally propose a model for the molecular mechanism behind the different behaviors of Arg/Trp rich CPPs (Fig. 10). In this model, peptides containing no or only one Trp behave as previously described in the literature. They act as positively charged polyelectrolytes interacting with negatively charged partners in a purely electrostatic manner. Their lack of membrane disruption [no cell membrane permeabilization, little effect on lipids in DSC experiments) suggests a non-disruptive translocation mechanism. Though our data shed no light on this particular aspect, we hypothesize it could be the so-called phase transfer mechanism, extensively described for such peptides [4–6]. Adding in more Trp (2 or 3) shifts or diversifies the behavior of the peptides. With the loss of 2 charges (Arg) and addition of 2 Trp, the peptides are less efficient. Most probably, the dominating effect is the loss of charges, as previously observed for polyarginines [53, 54]. On the other hand, when 3 Trp are present, these residues can compensate for the loss of charges. The hypothesis, supported by our ITC and DFT data, is that favorable interactions with GAGs are occurring thanks to the presence of Trp and point towards ion pair-π interactions. These interactions appear to be critical for GAG-dependent internalization. For direct translocation, our DSC data suggest that peptides containing 2 or 3 Trp promote lateral phase segregation, and this type of translocation mechanism has previously been proposed for CPPs [34, 41, 50]. Finally, with 4 Trp, internalization is greatly enhanced. However cytotoxic effects start to appear and thus, we suggest pores could be forming on the membrane, as a possible translocation mechanism [21, 62]. One point we would like to stress is that these behaviors are likely not mutually exclusive, but as adding more Trp, peptides gain new properties and diversify the way they can interact with their membrane partners, lipids and GAGs.

**Figure 10:**
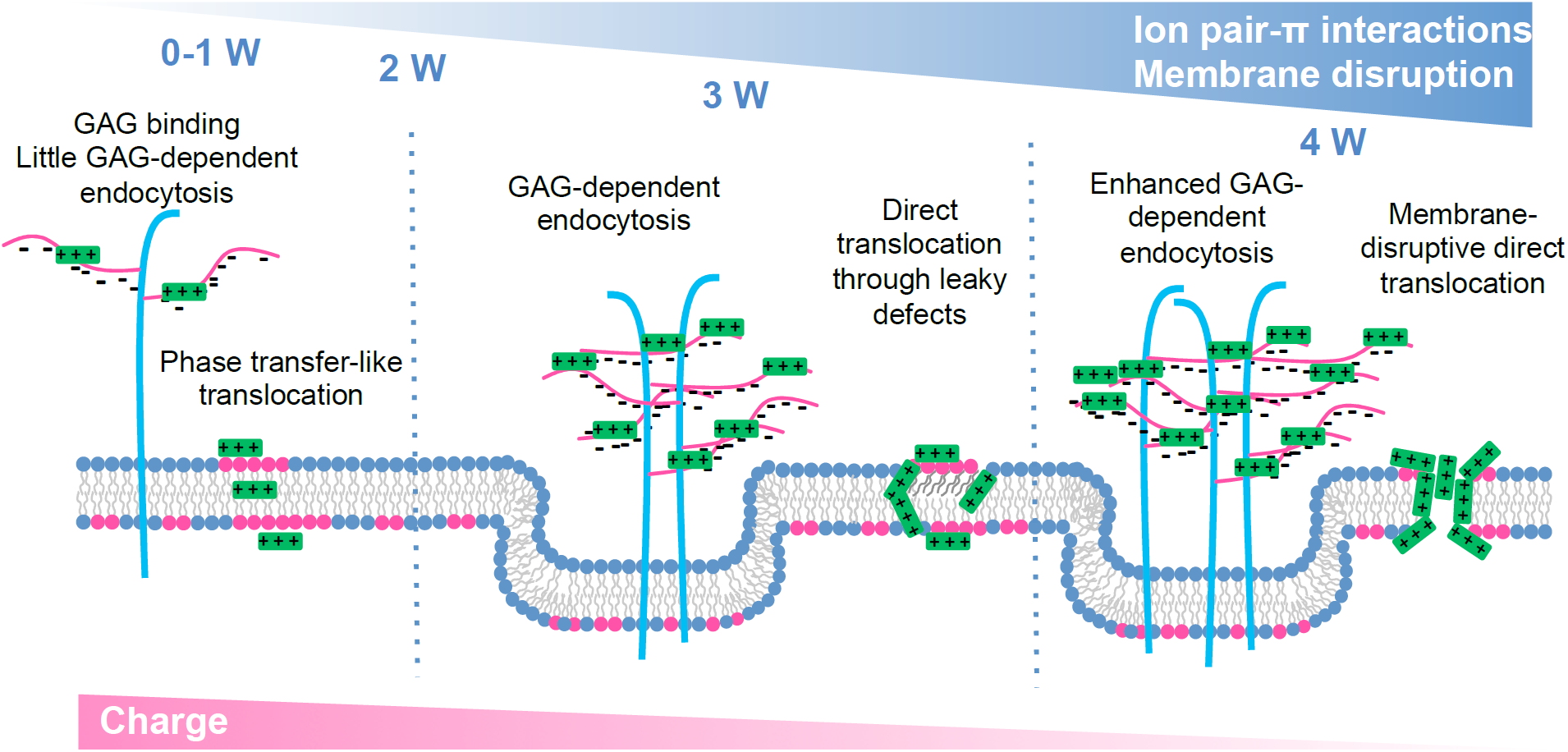
Proposed model for the internalization mechanisms of Arg/Trp peptides (see main text for detailed description).

## 5. Conclusion

The importance of Trp for some CPPs has been uncovered almost at the same time as CPPs themselves [7]. It has long been believed that hydrophobic residues coupled with a facial amphiphilic α-helical structure were an essential feature for some CPPs, probably by analogy with cationic antimicrobial peptides. At the same time, activation of Arg-rich CPPs with electron-rich aromatic systems such as pyrene butyrate has been used for more than a decade [63]. Herein, we show that facial amphiphilicity is not likely to play a crucial role for Arg-rich CPP uptake. Our work strongly highlights that Trp residues play the role of natural aromatic activators of Arg-rich CPPs, which happens through ionpair-π interactions. Beyond the field of CPPs, the existence of such energetically favorable ionpair-π interactions involving Arg, Trp and negatively charged moieties (carboxylate, sulfate, phosphate) could be of major importance for the comprehensive analysis of peptides or proteins (membranotropic antimicrobial, viral, and antitumor, receptor ligands etc.) interactions with lipids, polysaccharides or proteins.

## Supporting Information

Supporting Information include the detailed experimental procedures, mass spectra of peptides, helical wheel projections and structure simulations, ITC and IR data for all peptides (pdf file).

## Notes

The authors declare no competing financial interest

## Acknowledgments

We thank the Mass Spectrometry Platform (IBPS, Sorbonne Université) for services. This work was supported by the Agence Nationale pour la Recherche (ANR-10-BLAN-1417).

## Abbreviations

ART-FTIR: Attenuated Total Reflection Fourier Transformed Infra Red
CHO: Chinese Hamster Ovary
CPP: Cell-Penetrating Peptide
DFT: Density functional theory
DMPG: 1,2-dimyristoyl *sn*-glycero-3-phospho-(1’-rac**-** glycerol)
DSC: Differential Scanning Calorimetry
GAG: Glycosaminoglycan
ITC: Isothermal Titration Calorimetry
POPG: 1-palmitoyl-2-oleoyl-*sn*-glycero-3-phospho-(1’-rac**-**glycerol).

## Supporting Information

### Extended Experimental Methods

#### Peptide synthesis and purification

Boc-L-Arg(Tos), Boc-L-Trp(For), Boc-Gly, D-Biotin, MBHA Resin (0.53 mmol/g loading) and HBTU were purchased from Iris Biotech GmbH. Boc-(2,2-D_2_)-Gly was obtained from Cambridge Isotope Laboratories. Boc-D-Trp was purchased from Sigma-Aldrich.

D-Biotin was fully oxidized to D-Biotin sulfone (Biot(O_2_), Figure S1) by 4 days treatment with 30% H_2_O_2_ in H_2_O and used without further purification. This avoids further oxidation of the peptide throughout time.

**Figure S1:**
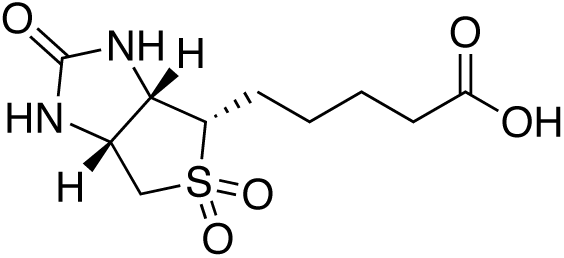
D-Biotin sulfone structure.

Peptides were synthetized by hand on a 0.1 mmol scale for non-deuterated peptides and 0.01 mmol for deuterated peptides. Amino acid (5 eq) activation was performed by HBTU (4.5 eq) in the presence of excess DIEA (12 eq), and Boc deprotection was performed in neat TFA (2× 1 min). Trp side chains were deprotected prior to cleavage by treatment with 10% piperidine in DMF (1, 2, 5, 10, 30 and 60 min successive incubations). Peptides were cleaved from the resin by anhydrous HF (2h, 0°C) in the presence of anisole and dimethylsulfide. Peptides were purified by reverse phase HPLC on a C18 preparative column (Macherey Nagel) with a H_2_O (0.1% TFA)/MeCN (0.1% TFA) elution gradient. Peptide purity and identity were further characterized by analytical reverse phase HPLC (C18, Higgins Analytical) with a H_2_O (0.1% TFA)/MeCN (0.1% TFA) elution gradient and MALDI-TOF MS (AB Sciex Voyager DE-PRO MALDI TOF or 4700 MALDI TOF/TOF)(Table S1, Figure S2).

**Table S1:**
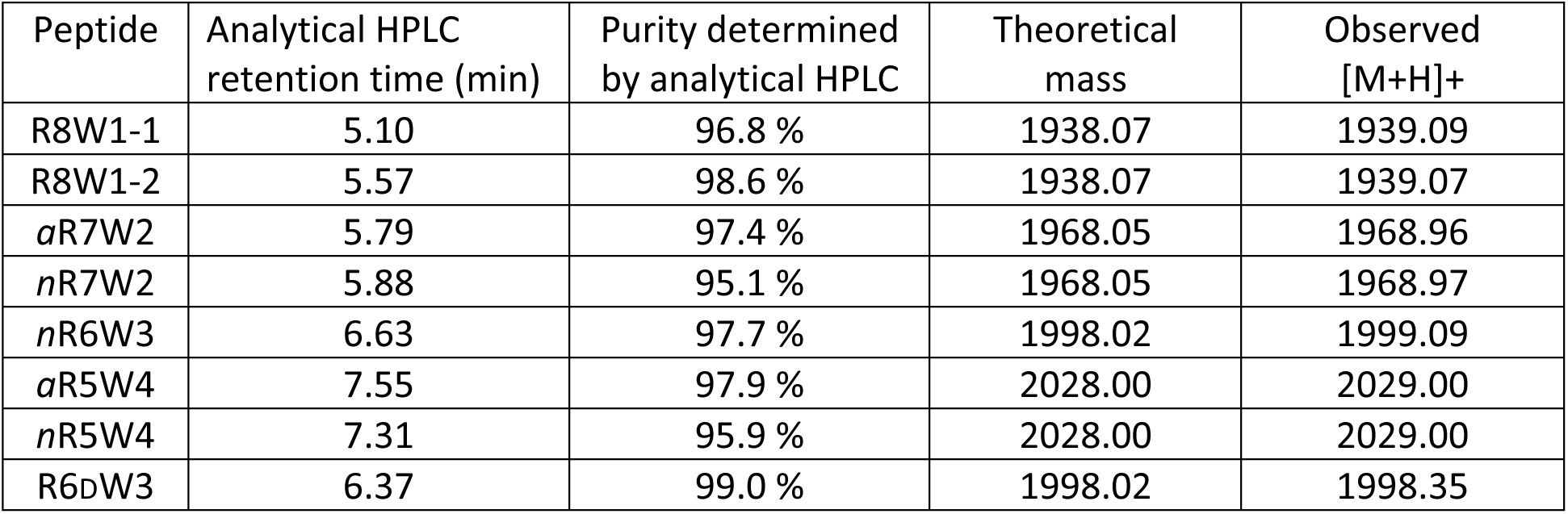
HPLC and MS characterization of the newly synthetized peptides. R9 and aR6W3 were previously synthesized and characterized (1).

**Figure S2:**
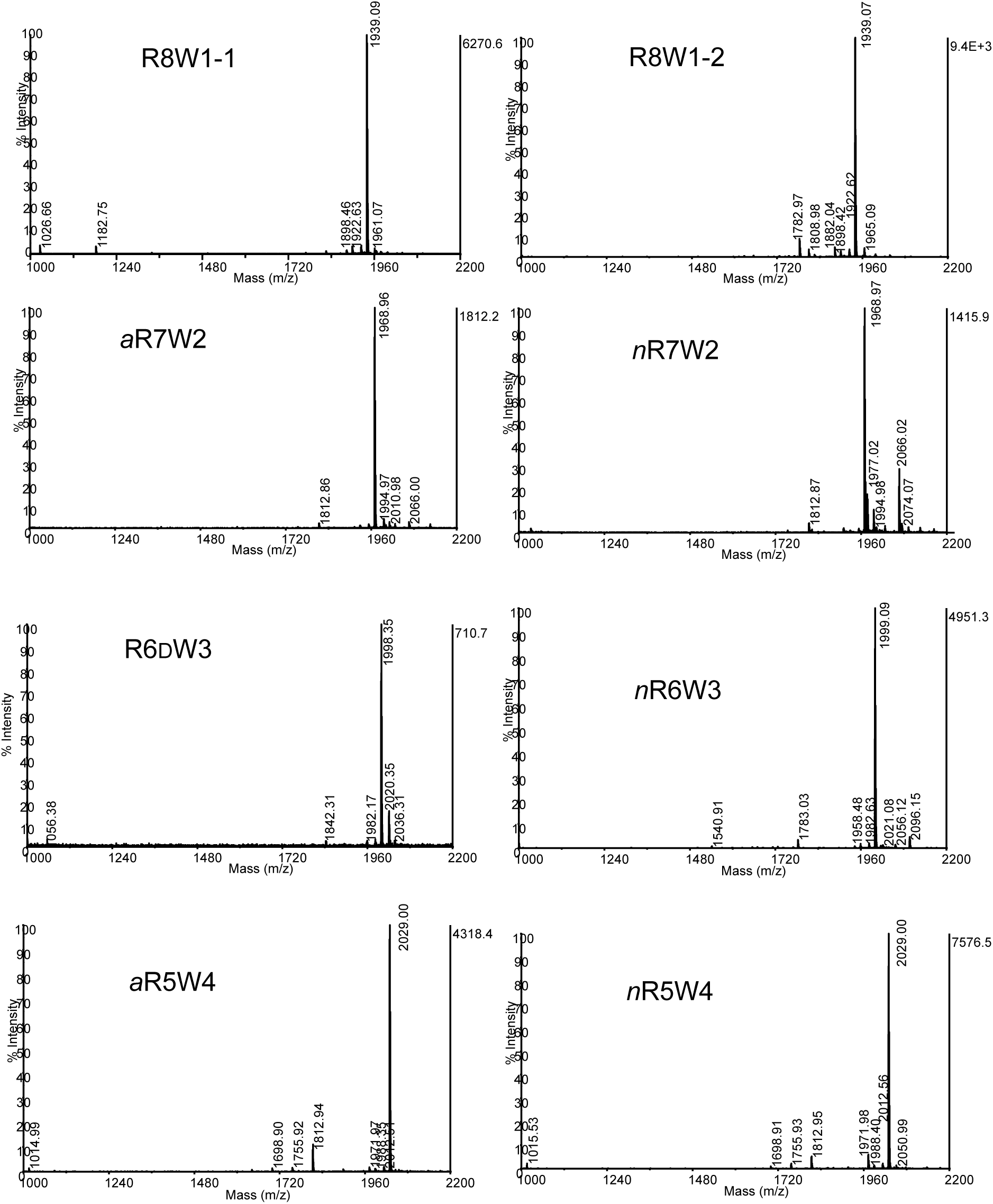
MALDI-TOF MS characterization of all the newly synthesized peptides.

### Extended Results

#### Peptide design

Helical wheel projections for all peptides are presented on figure S3.

**Figure S3:**
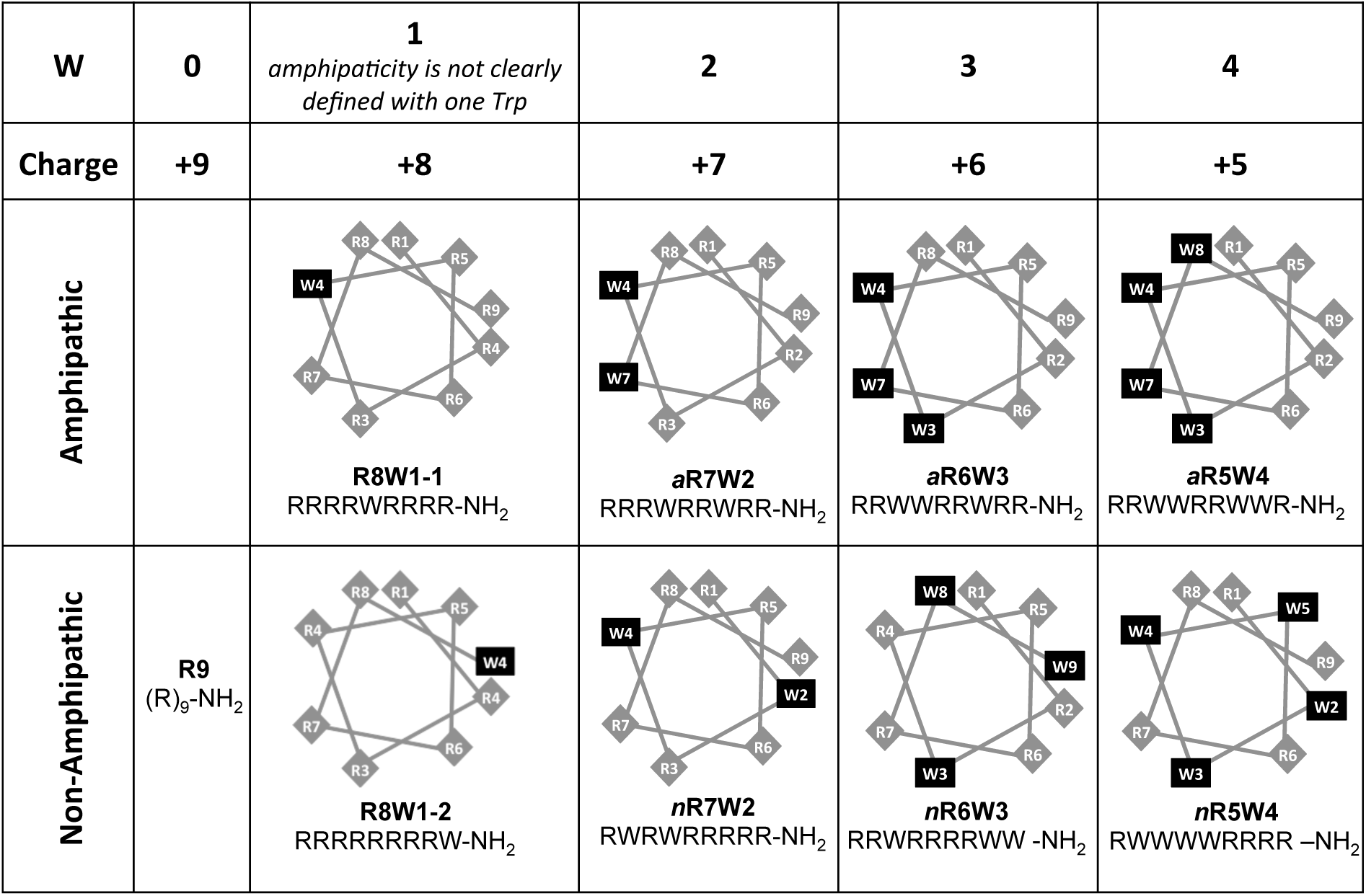
Helical wheel projections for all peptides.

#### Peptide binding to GAGs and lipids analyzed by ITC

All ITC experiments were performed with a NanoITC calorimeter (TA instruments) and data were analyzed using NanoAnalyze (TA instruments) using a simple binding model with *n* independent binding sites. The volume of the calorimetric cell is 983 μL and the injection syringe is 250 μL.

Representative ITC data injections of heparin (HI) into peptide solutions are shown on figure S4. Experiments were performed twice and the numbers given in tables 2 and 5 of the manuscript are averaged on the two experiments.

**Figure S4:**
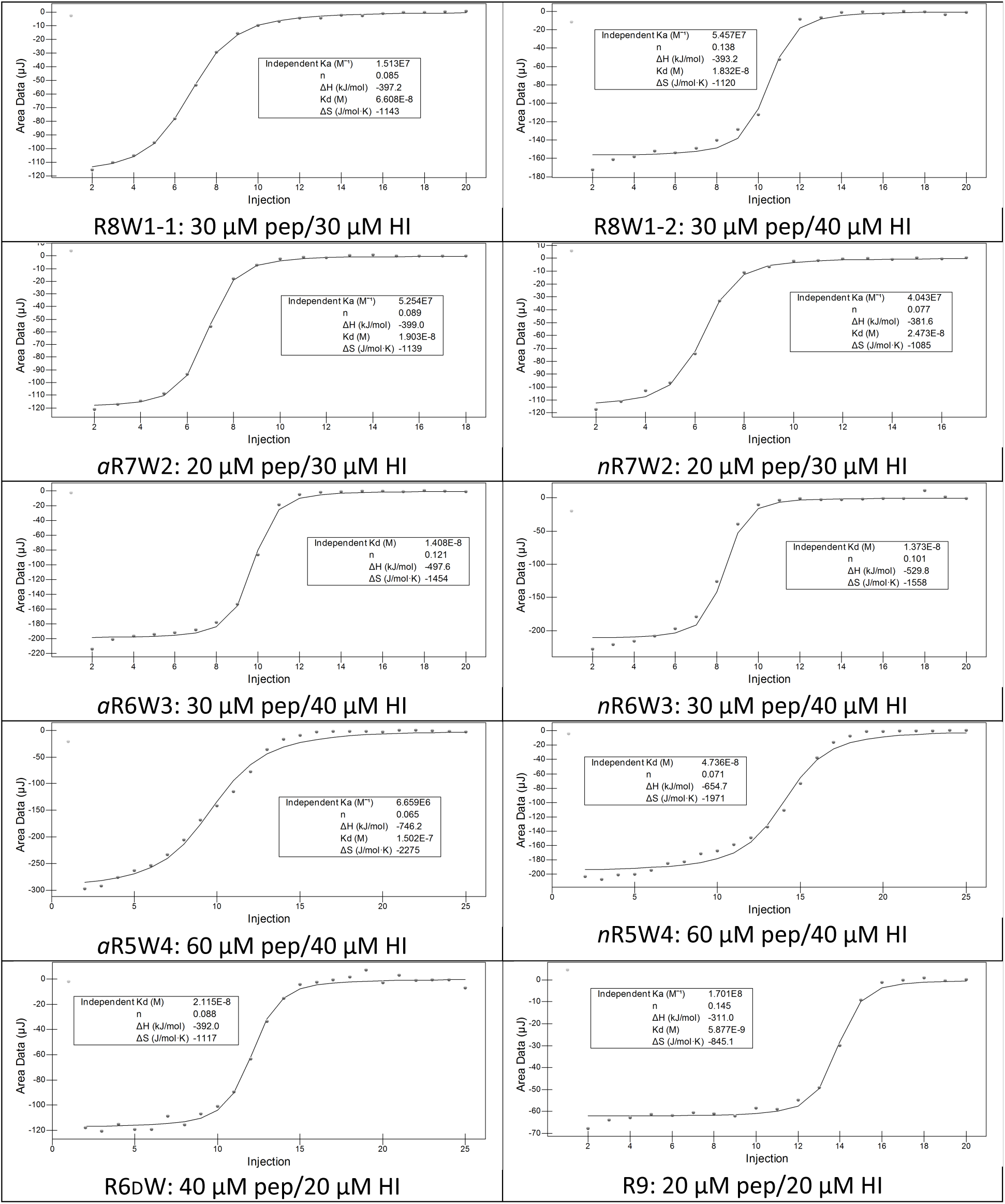
Binding isotherms for peptides binding to heparin as obtained after analysis by NanoAnalyze.

Representative ITC data injections of POPG LUVs into peptide solutions are shown on figure S4. Experiments were performed once or twice for each peptide.

**Figure S5:**
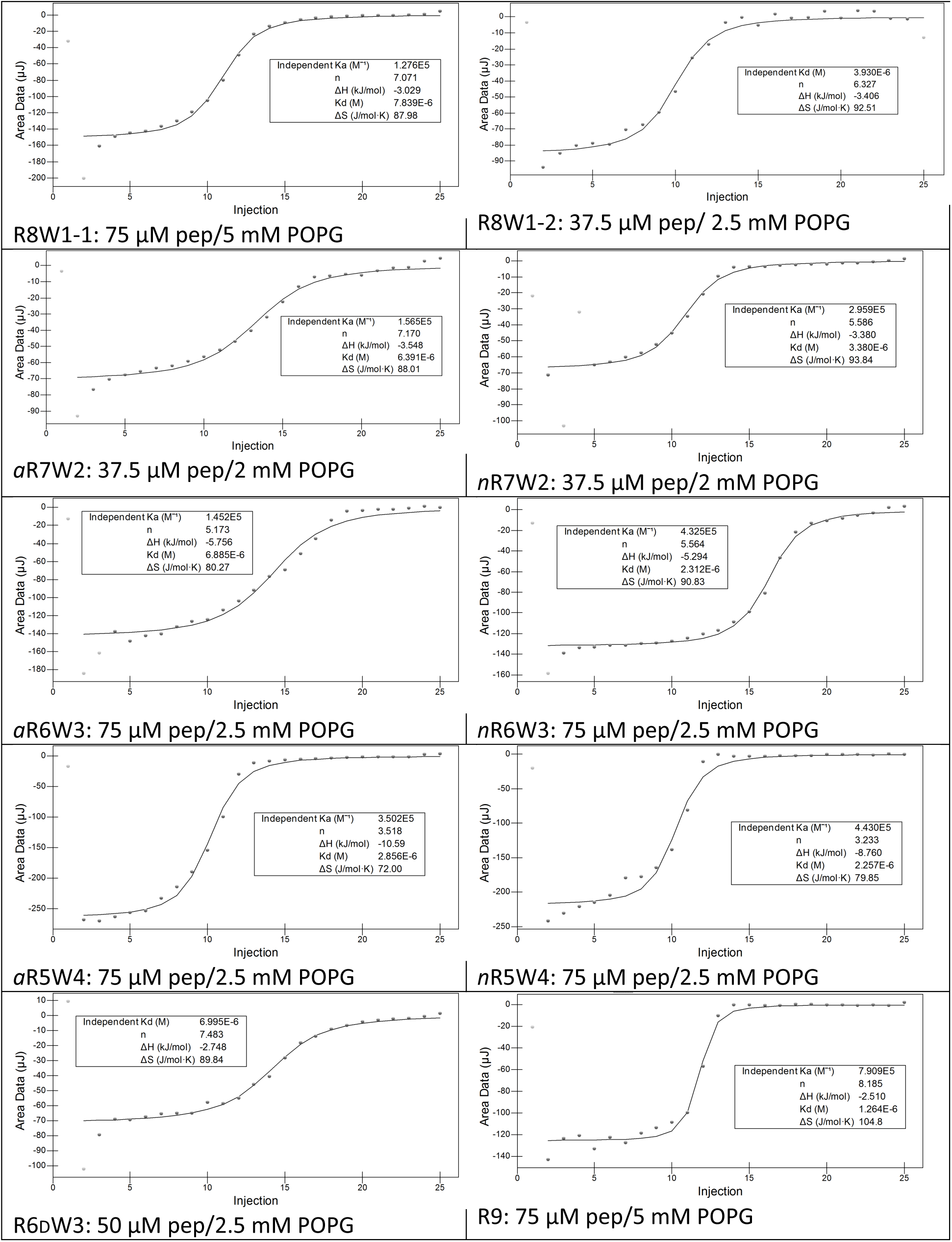
Binding isotherms for peptides binding to POPG LUVs as obtained after analysis by NanoAnalyze.

#### Peptide secondary structure

Example of Amide I band deconvolution for *a*R5W4 peptide is shown on figure S6 and secondary structures extracted from ATR spectra for all peptides are given in Table S2.

**Figure S6:**
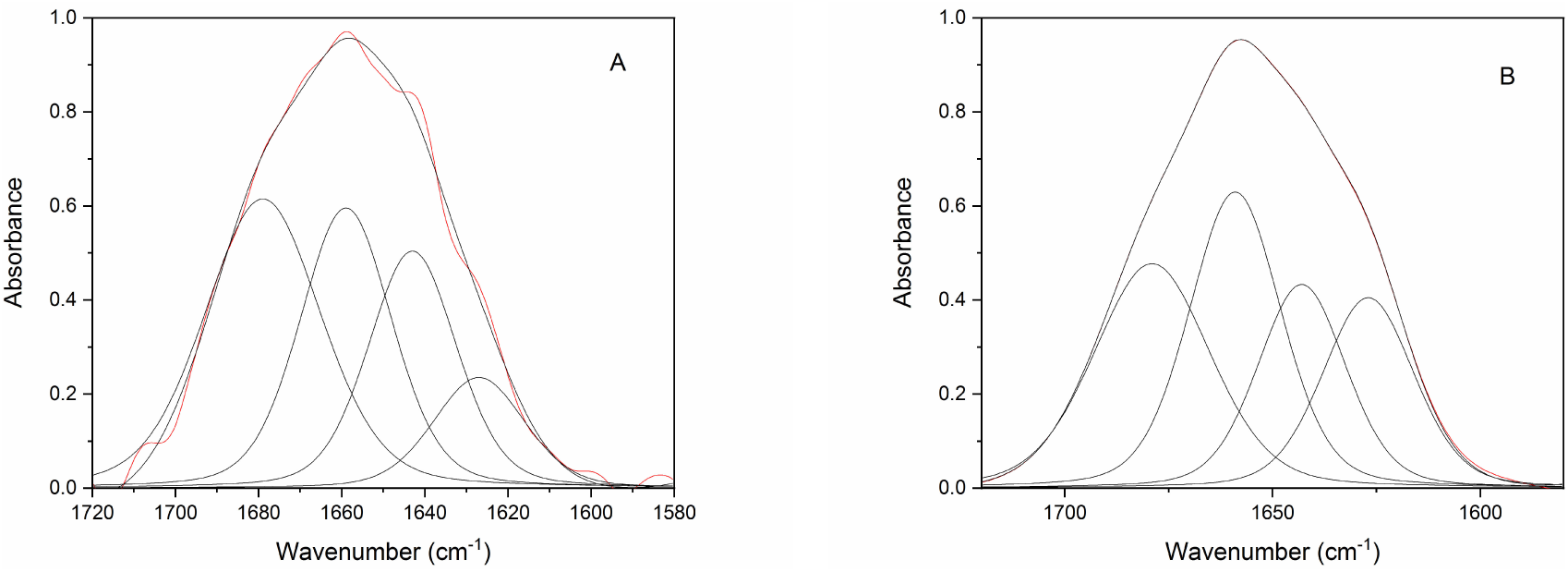
Example of ATR-IR spectra Amide I band deconvolution for *a*R5W4 alone (A) or with Hep (B)

**Table S2:**
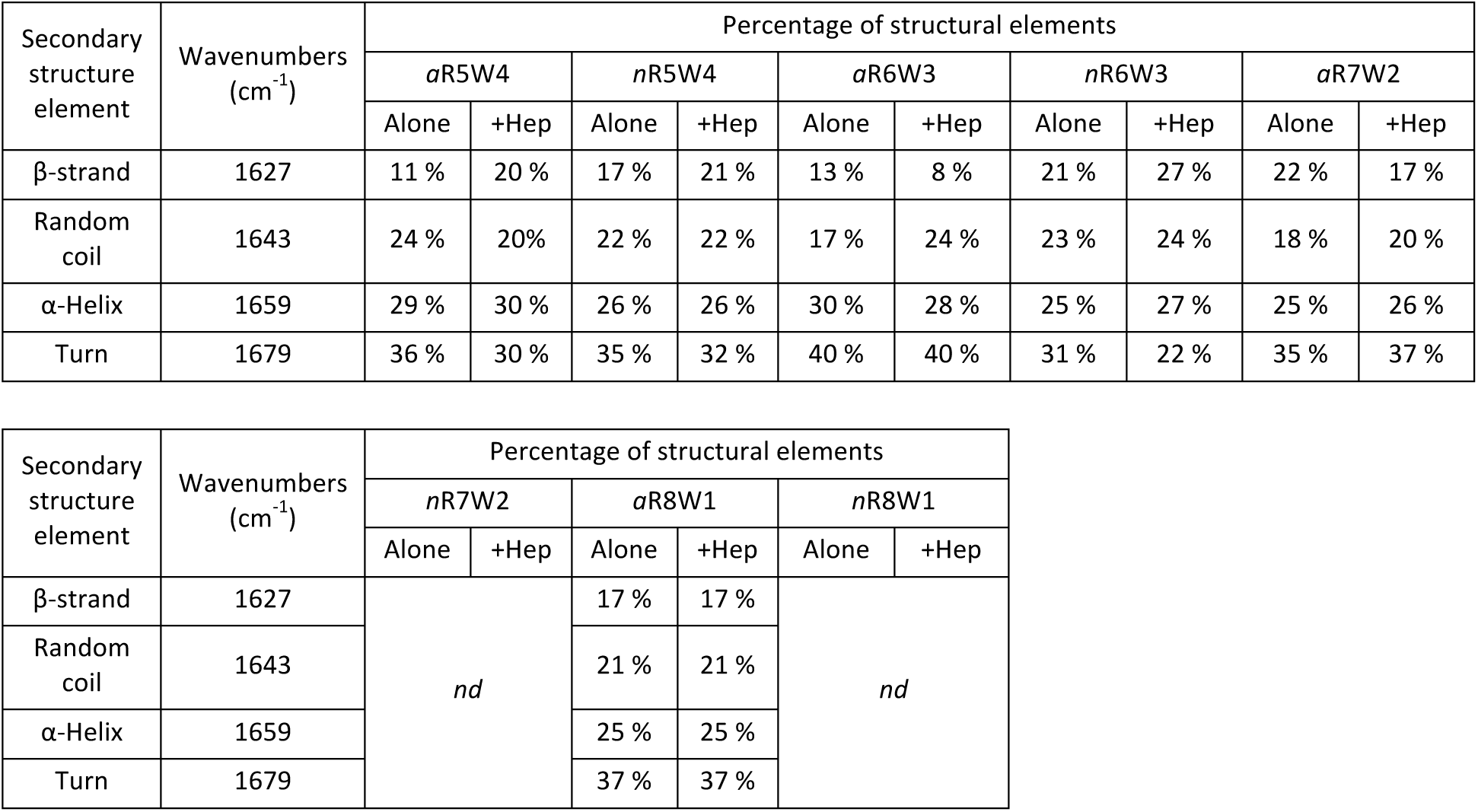
Secondary structure of all RW peptides in the absence or presence of heparin (8.3 μM heparin and peptide concentration according to the stoichiometries determined by ITC). For two peptides, secondary structures could not be extracted from the spectra. *nd*: not determined.

